# Entirely synthetic memory inception via dual optogenetic activation of sensory and dopaminergic neurons

**DOI:** 10.1101/2025.06.11.659083

**Authors:** Tayfun Tumkaya, Xianyuan Zhang, Yishan Mai, James Stewart, Adam Claridge-Chang

## Abstract

Associative conditioning is a fundamental learning paradigm that links salient stimuli to appropriate behavior. The vinegar fly is an important memory model that can associate odors with rewarding or punishing reinforcement stimuli via distinct sets of dopaminergic neurons. While much is known about learning, a crucial question remains: is the simple co-activation of sensory and neuromodulatory dopaminergic neurons sufficient to induce memory formation? To address this, we systematically replaced natural stimuli, namely odor-evoked activity, punishment, and reward signals, with optogenetic activation of corresponding neurons, both individually and in combination. In all cases, including a full-substitution paradigm where both olfactory activity and reinforcement dopamine signals were replaced, optogenetic activation successfully implanted synthetic memories. These findings provide a clear answer: simple coincident, rectangular-pulse sensory and neuromodulatory stimulation can instruct associative memory formation. We conclude that associative memory does not have strict requirements for temporal or ensemble activity patterns in olfactory or dopaminergic neurons that are specific to natural stimuli (e.g. odorants, sugar, or pain); rather, simple coincidence is an effective driver of memory inception. These findings open new avenues for manipulating and studying memory formation with all-optical control.

## Introduction

Many animal species establish connections between two different stimuli that occur at or around the same time via a process known as associative learning. Using conditioning methods to induce associative learning, scientists have elucidated many molecular and circuit properties of memory. Associative-conditioning paradigms typically use an unconditioned stimulus (US) to train a conditioned stimulus (CS). In this way, various forms of US–CS associative sensory learning have been demonstrated in the vinegar fly *Drosophila melanogaster*, including visual, gustatory and place learning. Most research has focused on olfactory conditioning (Owald and Waddell 2015; Cognigni, Felsenberg, and Waddell 2018; Amin and Lin 2019; Modi, Shuai, and Turner 2020; Davis 2023), which depends on the contingency of odor and reinforcement signals. For example, flies can associate odors (CS) with either rewarding or aversive stimuli (US), such as sugar or electric shocks, respectively.

Odorants are bound and detected by olfactory receptor neurons (ORNs) that communicate via projection neurons (PNs) to higher brain centers, namely the lateral horn (LH) and the mushroom body (MB) (Masse and Turner 2009; Benton 2022; Choi, Kim, and Hyeon 2022; Fulton et al. 2024). The MB is a crucial site where synaptic plasticity occurs, which is necessary for memory formation (Aso et al. 2014; Cohn, Morantte, and Ruta 2015; Hancock et al. 2022). Rewarding and aversive reinforcement signals are transmitted to the MB via distinct sets of dopaminergic neurons: aversion signals are mediated by the paired posterior lateral 1 (PPL1) neurons, whereas reward signals are conveyed by the paired anterior medial (PAM) neurons (Schwaerzel et al. 2003; Claridge-Chang et al. 2009a; Burke et al. 2012; C. Liu et al. 2012). Following a conditioning paradigm, coincident or temporally contingent activity in PPL1 or PAM neurons with the olfactory pathway was shown to lead to changes in synaptic strength between odor-activated MB neurons and their output neurons (Owald and Waddell 2015; Modi, Shuai, and Turner 2020). Many such findings have been made using interventions that silenced or activated specific neuronal types.

Activation has historically been used to determine which neuronal signals can induce associative conditioning. Here, a US or CS is replaced with experimentally induced activity in a substitution experiment. Pioneering *in vitro* work demonstrated that the neurotransmitter serotonin (5-HT) is sufficient to establish synaptic plasticity between *Aplysia* sensory neurons and gill motor neurons (Montarolo et al. 1986; Rayport and Schacher 1986). Meanwhile studies performed in mice have investigated the synaptic model of learning by labeling neurons that are active during memory conditioning with opsins. These activity-labeled ‘engram’ neurons have subsequently been optogenetically manipulated to induce memory retrieval and association with novel memories (X. Liu et al. 2012; Ramirez et al. 2013; Yiu et al. 2014; Rashid et al. 2016).

In *Drosophila*, the olfactory conditioning paradigm has been reconstituted by replacing either the sensory or neuromodulatory stimulus with artificial neural activation, using optogenetic and/or thermogenetic neural activation approaches. On the sensory (CS) side, artificial ORN stimulation effectively substituted for genuine odors in olfactory conditioning (Tomasiunaite, Widmann, and Thum 2018). On the neuromodulatory (US) side, synthetic dopaminergic PAM or PPL1 neuron activation during exposure to authentic odors led to the formation of appetitive and aversive olfactory memories, respectively (Schroll et al. 2006; Claridge-Chang et al. 2009a; Aso et al. 2010; C. Liu et al. 2012; Burke et al. 2012; Aso et al. 2012; Dawydow et al. 2014; Hige et al. 2015; Aso and Rubin 2016). Dopaminergic activation can also induce gustatory learning (Jelen et al. 2023). Both PAM neurons and their upstream P1 neurons, both implicated in male courtship behaviors, imparted positive valence to otherwise neutral odors in male flies when stimulated optogenetically (Shen et al. 2023).

Most synthetic-memory experiments have used a one-pathway design where activity is only actuated in either CS- or US-related neurons. At least two studies have employed completely artificial neuronal activity in a two-pathway design (Honda et al. 2014; Vetere et al. 2019). One study performed in *Drosophila* larvae substituted both the odor and reward signals by artificially and concurrently activating ORNs and Tyrosine decarboxylase 2 (Tdc2) cells (upstream of the dopaminergic PAM neurons); the result was the induction of an entirely fictive appetitive memory (Honda et al. 2014). The other study was performed in mice, whereby concurrent optogenetic stimulation of a single olfactory glomerulus and either the laterodorsal tegmentum (LDT) or the lateral habenula (LHb) neurons (which synapse onto dopaminergic neuromodulatory circuits) could artificially implant attractive or aversive memories, respectively (Vetere et al. 2019). Notably, while these two-pathway memory implantation studies showed that neuromodulatory signals can be substituted with artificial activation of the upstream neurons, the instructive dopaminergic circuits themselves were not directly engaged.

Several neural-system properties can occur in natural conditioning and might persist even in two-pathway memory studies as the artificial US signal is relayed to the downstream dopaminergic neurons. Examples of such properties could include temporal dynamics (Cassenaer and Laurent 2012), specific cellular ensembles (Takemura et al. 2017; Zhao et al. 2021), or pathways activated beyond the major identified circuits (Zeng et al. 2023).

Inception of entirely synthetic memories via optogenetic actuation would support the idea that straightforward activation of the targeted pathways is sufficient to instruct associative learning. Thus, an unresolved question remains: would the simple co-activation of sensory cells and the plasticity-inducing dopaminergic neurons successfully implant memory?

Here, we aimed to answer this question using olfactory conditioning in adult *Drosophila* to assess the olfactory–dopamine convergence model by directly activating both of the neurons that transmit odor and reinforcement signals. We hypothesized that by simultaneously actuating the CS and US pathways, both aversive and attractive olfactory-like memories could be implanted. To this end, we developed an entirely artificial conditioning paradigm using two opsins with distinct spectral properties (red-shifted Chrimson and blue-shifted Chr2XXL) to independently manipulate the ORNs and transgenic drivers that included the PAM or PPL1 neurons. The experimental design included: (1) an absence of fine temporal structure in the stimulation schedule beyond simple on–off illumination; (2) almost no spatial or ensemble structure in the ORNs, but instead, a broad activation of 70% of all receptor cells; and (3) very limited ensemble structure in the neuromodulatory cells, targeting broad expression either in the aversive or appetitive dopaminergic neurons (Wang et al. 2003; Larsson et al. 2004). This approach allowed us to investigate whether fully synthetic conditioning paradigms could enable robust artificial memory inception.

## Material and methods

### Fly care and strains

Flies were reared on standard cornmeal-based media at 25°C degrees, under a 12 h light : 12 h dark cycle, except for the flies used in optogenetic experiments. For optogenetic experiments, flies were kept in the dark throughout their development and transferred to 0.5 mM all-trans-retinal (ATR; Sigma-Aldrich, USA)-supplemented food 2 days before the experiments. The wild type genotype used was cantonized *w^1118^*. Transgenic *Orco-Gal4* RRID:BDSC_26818, BDSC_23292 (Larsson et al. 2004; Ray et al. 2007), *TH-LexA* RRID:BDSC_99050 (Berry et al. 2015), and *UAS-Chrimson* RRID:BDSC_55135 (Klapoetke et al. 2014) files were obtained from the Bloomington *Drosophila* Stock Center. The *LexAop-Chr2XXL* stock was a gift from Christian Wegener (University of Wurzburg) (Selcho et al. 2017).

### Multi-fly olfactory trainer (MOT) assay

The MOT assay has been described previously (Claridge-Chang et al. 2009a; Mohammad et al. 2017; Kan et al. 2021). Briefly, it is conducted using an apparatus comprising eight chambers (50 × 5 × 1.3 mm) with acrylic walls, a glass floor and ceilings printed with transparent indium tin oxide electrodes (ITO; Visiontek UK). The transparent ITO boards deliver a foot shock to the flies, while the transparency permits fly tracking (up to 8 flies per chamber) using a guppy camera (AVT F-080) and in-house software, CRITTA, an instrumentation and image-processing software written in LabVIEW (National Instruments). The chambers have two odor inlets on each end and two odor outlets in the middle. The odors 3-octanol (OCT) and 4-methylcyclohexanol (MCH) flow into the chambers via the odor inlets; the odor concentrations are then adjusted using mass flow controllers (MFCs; Sensirion AG, Sweden). For classical MOT conditioning experiments, 6–8 flies were ice-anesthetized and transferred into the MOT chambers. The conventional training protocol was as follows: after a brief 1-minute acclimatization, flies were trained by 12, 60V foot shocks along with the first odor for 1 min before a second 1-min acclimatization and exposure to the second odor for 1 min without a foot shock. After the training session, the flies were gently pushed to the middle of the chamber (choice point) using air puffs from both ends after 1 min of compensatory air, and tested between the two odors for 2 min, as follows:

1. Bias Check, 2 min
2. Wait, 1 min
3. Train 1, 1 min
4. Wait, 1 min
5. Train 2, 1 min
6. Wait, 1 min
7. Puff
8. Test, 2 min

For non-odor optogenetic conditioning, Train 2 was omitted, and rather, the single Train epoch was followed by 3 min of Wait prior to the Test. The Bias Check epoch was used to establish the baseline preference of the flies, which may vary due to technical factors (e.g., light effects) or biological factors (e.g., innate responses triggered by the target ORNs) (Claridge-Chang et al. 2009b).

### Wind- and light-induced self-administration response (WALISAR)

WALISAR was used to assess individual fly behavior, as opposed to groups of flies (6-8) per chamber (in the MOT assay). The apparatus was set-up as previously described (Tumkaya et al. 2022), using blue and red LEDs. Briefly, the chambers were formed from rectangular acrylic assemblies (11.5 × 14.5 × 0.3 cm), each of which contained 26 disco-rectangular chambers (50 × 4 × 3 mm) with airflow inlets and outlets at each end. A stopcock valve (Cole-Parmer) was used to modulate the flow rate of compressed air to give a chamber wind speed of ∼35 cm/s, which was verified using an airflow meter (Cole-Parmer).

Optogenetic light was delivered with LEDs [LUXEON Rebel LEDs on a SinkPAD-II 10 mm Square Base; red (617 nm), green (530 nm), blue (470 nm), each equipped a lens (17.7° 10 mm Circular Beam Optic)] and positioned above the chamber assembly at a 45° angle. CRITTA software was used to automate the timing of the LEDs throughout each experiment. The stimuli exposure schedule was similar to that used during the MOT assay, with a few differences: no air puff was given prior to the Test, and air flowed from one end to the other rather than using a convergent flow. The resulting protocol was as follows:

1. Bias Check, 2 min
2. Wait, 1 min
3. Train 1, 1 min
4. Wait, 3 min
5. Puff
6. Test, 2 min

### Optogenetic substitution experiments

Optogenetic substitution experiments used the same durations per training step as the conventional olfactory conditioning protocol in the MOT assay. In the odor substitution experiments, a PC-controlled LED micro-projector (Optoma ML750) was used to activate the Chrimson channel and to generate a choice pattern by illuminating half of the arena with red light. Blue LEDs (Luxeon Rebel LEDs on a SinkPAD-II 10-mm Square Base available from http://www.luxeonstar.com; blue SP-05-B4, peak 460 nm) were placed above the arena and served to replace the foot shocks with blue-light shifted Chr2XXL channelrhodopsin. For aversive memory implantation, 12 light pulses were delivered in the same temporal pattern as used during standard olfactory conditioning (1.5 s ON – 4.5 s OFF). For reward memory implantation experiments, the blue light was turned on only for the first 3 s of the 60-s training session as Chr2XXL is slow to close (Dawydow et al. 2014).

### Immunohistochemistry

*Drosophila* brains were dissected in cold phosphate buffered saline (PBS) and fixed in 4% paraformaldehyde (Electron Microscopy Sciences) in PBT (a mixture of PBS and 0.3% Triton X-100) for 20 min. Then, the samples were washed three times with PBT and blocked with 5% normal goat serum in PBT for 1 h. Primary antibodies were applied to the samples and left to incubate overnight at 4°C. The samples were washed with PBT three times (10 min each) and incubated with secondary antibodies overnight at 4°C in the dark. The following primary and secondary antibodies were used in the procedure:

● Chicken anti-GFP antibody (ab13970, 1:1000),
● Mouse anti-Channelrhodopsin 2 (PROGEN, Cat # 651180, 1: 250),
● Alexa Fluor 568 goat anti-mouse (Invitrogen, Cat # A-11004, 1:500),
● Alexa Fluor 488 goat anti-chicken (Invitrogen, Cat # A-11039, 1:500).

Confocal images of the brain samples were captured under an LSM710 Carl Zeiss confocal microscope. The maximum-intensity projections of the confocal images were generated in ImageJ (Schneider, Rasband, and Eliceiri 2012; Schindelin et al. 2012) and the colors were inverted using the EZreverse web application (Song and Goedhart 2024).

### Data analysis

Throughout the MOT and WALISAR experiments, the coordinates of each fly were logged into .csv files via CRITTA. These data were then used to calculate the odor preference of the flies with custom Python scripts. Specifically, the number of flies in each odor zone was counted over the last 30 s of the test session and then reduced into one number per odor zone by averaging (Kan et al. 2021). The average number of flies in the unconditioned odor was then subtracted from the number of flies in the conditioned odor, and divided by the total number of flies, resulting in a chamber preference index (PI). Instead of full PIs, as is done conventionally (Tully and Quinn 1985), this study reports and plots half PIs, i.e. PIs for each anti-MCH or anti-OCT chamber. The results are presented as scatter plots, in which each dot represents one experiment consisting of 6–8 flies. The Bias Check, an internal control in each experiment, was used as the baseline preference of the flies. The change in preference (ΔPI) between an initial Bias Check and the Test was used as the learning effect size (Claridge-Chang et al. 2009b), calculated as the mean difference between the animals in their naive and conditioned states as in the formula below; the error bars represent bootstrapped 95% confidence intervals (CIs).

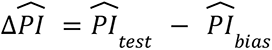

where 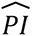 denotes the mean PI for test or bias group.

## Results

### Optogenetic fly lines successfully learn to avoid shock-paired odors

In our first experiment, we wanted to confirm that transgenic flies expressing the Chrimson opsin in their ORNs (*Orco>Chrimson* flies) were capable of normal olfactory learning. To do so, we used an olfactory classical-conditioning apparatus, referred to as the Multifly Olfactory Trainer (MOT) assay, to facilitate the automated training and tracking of small groups of flies (Mohammad et al. 2017; Kan et al. 2021). The training protocol included the presentation of the first odor with a foot shock, then a second odor without a foot shock, and finally a test session presenting a choice between the two odors (see Methods, **Figure 1A-C**). We tested three transgenic fly lines carrying the transgenes required for optogenetic activation of the *Orco-Gal4* neurons to verify normal shock learning (**Figure 1A**). We used tracking data to measure the flies’ odor preference before and after conditioning, and then calculated the change in preference (ΔPI) as a learning score (Claridge-Chang et al. 2009a). Here, we found that all three transgenic lines avoided the shock-paired odors: *UAS-Chrimson* ΔPI = −0.61 [-0.97, −0.33]*; Orco-Gal4* ΔPI = −0.72 [-0.90, −0.45]; and *UAS-Chrimson; Orco-Gal4* ΔPI = −0.44 [-0.84, −0.01] (**Figure 1D**). We found differences in the baseline preferences (PI) and the learning scores (ΔPI) between the three lines, with an unweighted average of the three ΔPIs of –0.59. These findings establish that these transgenic fly lines can learn.

**Figure 1.**
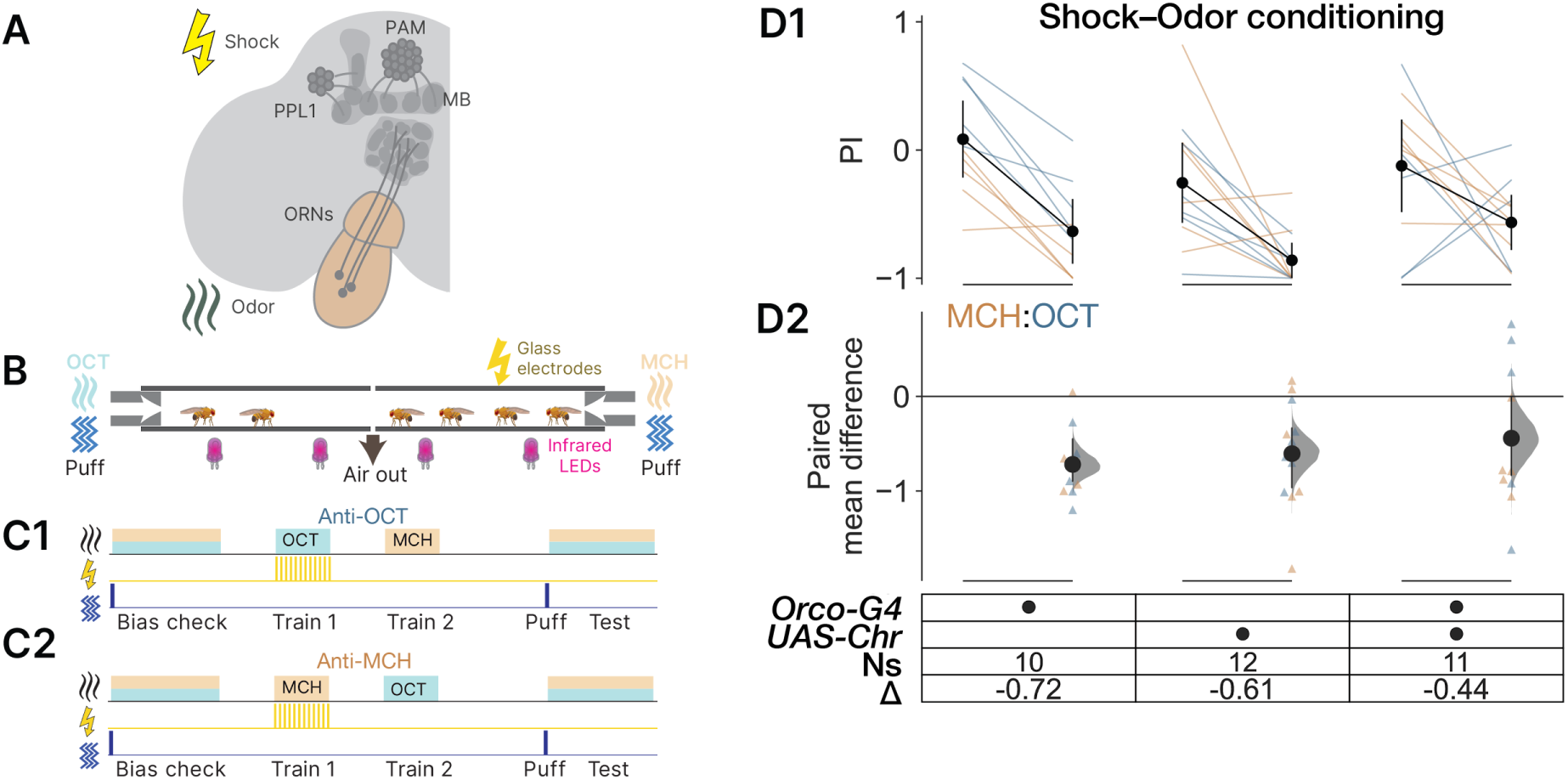
Shock-paired conditioning induces memories with real odors. **A.** Schematic of the key brain regions involved in olfactory learning: a mushroom body (MB), paired-anterior-medial (PAM) neurons, paired-posterior-lateral 1 (PPL1) neurons, the antenna and ORNs are shown. The shock and odor icons indicate the stimuli used in the experiment. **B.** Schematic of the MOT chamber showing odors (OCT and MCH), air flow with air puffs, infrared LEDs for imaging, and glass electrodes for shock. **C1.** The anti-OCT training paradigm: Flies (n=6-8) were loaded into chambers and tested for OCT and MCH preference (Bias check), then exposed to OCT with a foot shock (Train 1), MCH exposure without a shock (Train 2), agitated with an air puff, and finally re-tested for odor preference (Test). **C2.** The anti-MCH training protocol: as for the paradigm in C1, except that MCH was paired with a foot shock (Train 1). **D1.** Paired plot showing the chamber preference index (PI) before and after conditioning. The ends of the orange and blue lines indicate the PI scores per chamber for MCH and OCT training, respectively; black dots indicate the mean PI, and the error bars represent 95% CIs. **D2.** Change in chamber preference index (ΔPI) between pre- and post-conditioning is shown. The orange and blue triangles indicate the individual pre:post ΔPI values; black dots indicate mean ΔPI values with their 95% CI (black line), and the Δ distributions (gray curve).

### Optogenetic activation of Orco neurons can substitute for odor stimulus in conditioning

Having shown that *Orco>Chrimson* flies could learn a shock+odor association, we next wanted to test whether actuation of Chrimson could substitute for natural odors. To do so, we replaced odorants with the optogenetic activation of Orco ORNs using the red-shifted channelrhodopsin Chrimson during the shock epoch (**Figure 2A–C**). We then used the flies’ preference for red light before and after conditioning to calculate the ΔPI learning score. We saw that the optogenetically sensitive *UAS-Chrimson; Orco-Gal4* flies exhibited a strong aversion to red-light stimulation after conditioning, as evidenced by a ΔPI of −0.92 [-1.35, −0.44], while the light preference of the control flies shifted less (*UAS-Chrimson*, ΔPI = −0.21 [-0.57, +0.19]; *Orco-Gal4*, ΔPI = −0.07 [-0.49, +0.43], **Figure 2D**). While it is possible that some of the conditioning effect we saw was due to visual learning (Vogt et al. 2014), the large difference between the controls and the optogenetic scores (−0.07, −0.21 ≉ −0.92) indicates that visual learning is a minor contributor at most. Overall, this experiment established that optogenetic Orco neuron activation can replace an odor stimulus and serve as the substrate of aversive shock conditioning.

**Figure 2.**
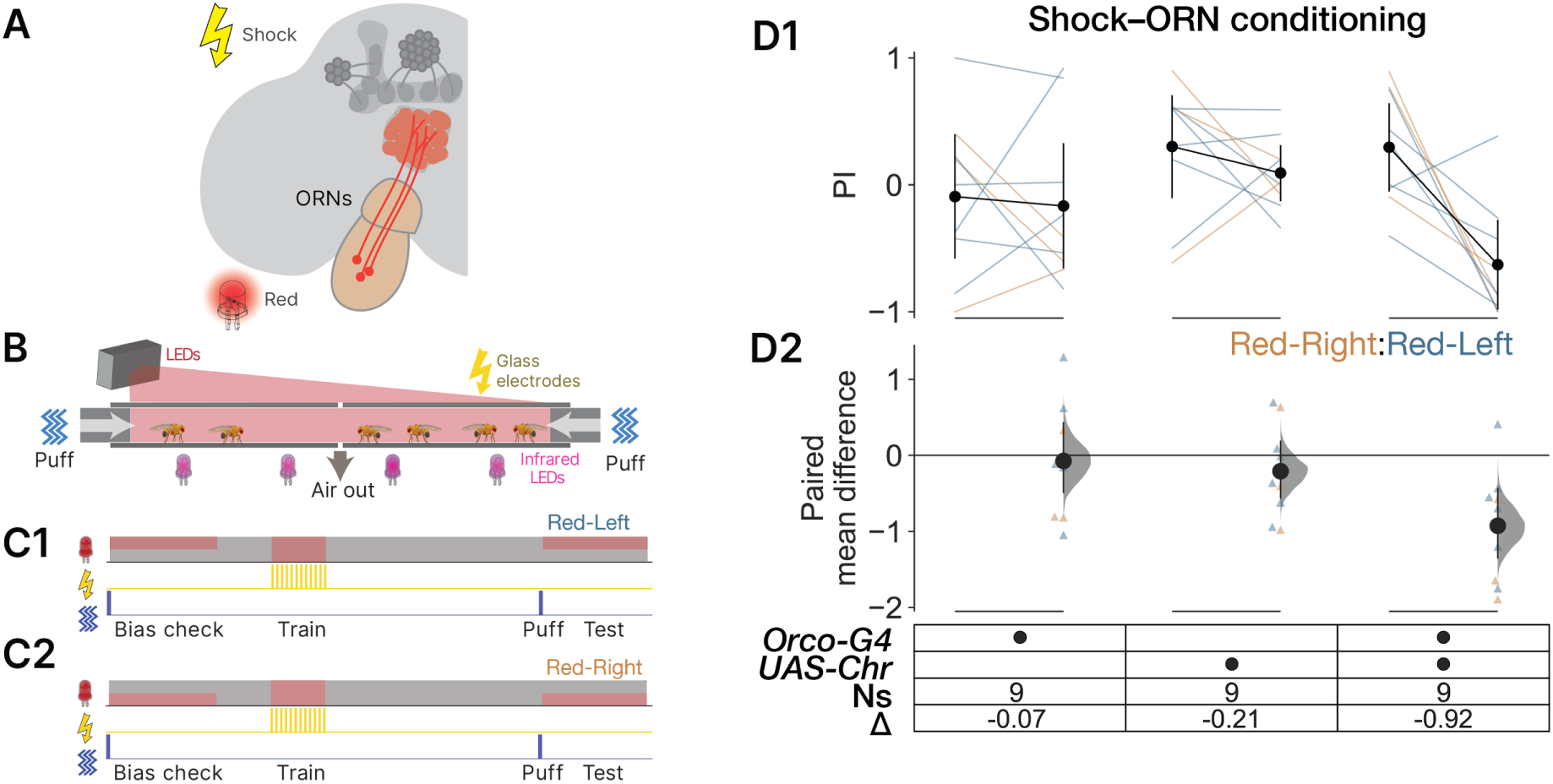
Shock-paired conditioning induces memories with a fictive odor-like stimulus. **A.** Schematic of the stimuli and targeted brain regions used for shock conditioning of a synthetic optogenetic olfactory-like stimulus. **B.** Schematic of the odor-free semi-artificial conditioning experiment, where Chrimson is activated with a red LED. **C1.** The Red-Left odor-substitution conditioning protocol: flies were tested between red light and no light (Bias), then trained with red light paired with a foot shock (Conditioning), a no-light interval, and finally a second light–dark choice task (Test). **C2.** The Red-Right odor-substitution conditioning experiment. The protocol was as for the Red-Left experiment in G1, but the light pattern was swapped. **D1.** Bias and Test PI scores for three genotypes, before and after training. The ends of each line indicate scores from one chamber (6-8 flies). Orange and blue indicate the Red-Right and Red-Left illumination schedules. Error bars are 95% CIs. **D2.** The relative ΔPI after conditioning is shown. The orange and blue triangles represent the individual pre:post ΔPI values per chamber where either a Red-Right or Red-Left light pattern was used, respectively. The black dots indicate the mean ΔPI for each group with its 95% CI (black line), and the Δ distribution (gray curve).

### Temporal structure of optogenetic stimulus is untrainable

Temporal patterns of ORN spikes may play a role in encoding odor identity and olfactory memory formation (Wilson, Turner, and Laurent 2004; Gruntman and Turner 2013; Groschner and Miesenböck 2019; Benton 2022). To investigate whether the temporal structure of an odor-like optogenetic stimulus carries salient information for conditioning (Tumkaya et al. 2019), we designed a series of experiments using fictive opto-odors. Flies were exposed to two types of fictive opto-odors (CS) by optogenetically activating their ORNs using distinct light-delivery protocols: (1) static red-light stimulation (“static odor”) and (2) pulsed red-light stimulation at a frequency of 20 Hz (“pulsed odor”). During training, either the static or pulsed odor was paired with foot shocks (US), while the other fictive odor was presented without shocks. In the test phase, flies were evaluated on their ability to avoid the punished optogenetic temporal pattern.

We reasoned that if the temporal pattern of the optogenetic light stimulus was a perceptible and learnable characteristic, flies would form an aversion specific to the punished temporal pattern. However, across the three types of ORNs tested in this paradigm (*Orco*, *Gr21a*, and *Or42b*), flies generally failed to learn the punished stimulus (**Figure 3**). The only exception was conditioning of anti–static-light activation of the *Or42b* neurons (**Figure 3C,D**). These findings suggest that precise temporal coding of ORN activity is not a critical determinant for olfactory perception or associative learning in these neurons.

**Figure 3.**
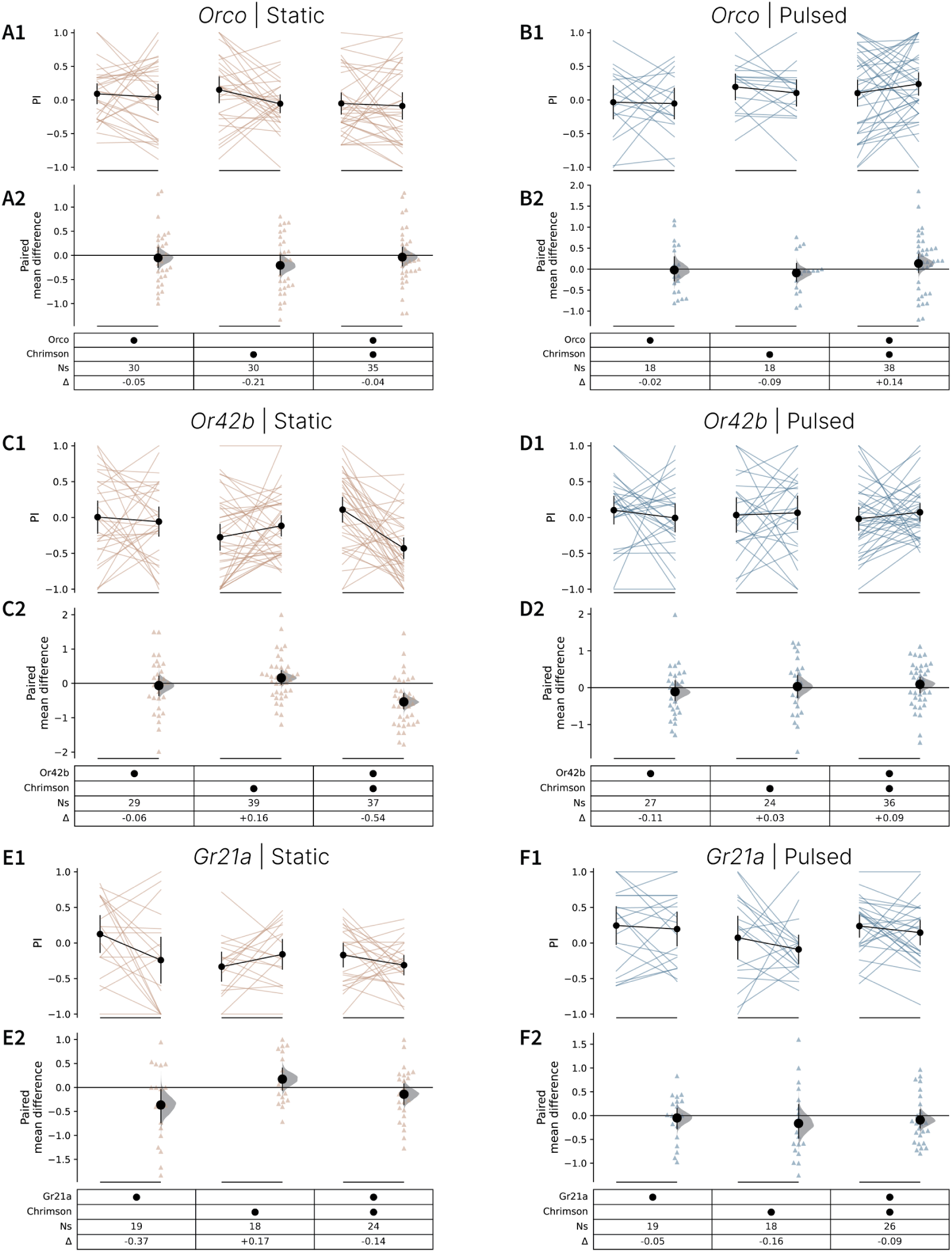
Temporal structure of the optogenetic stimulus targeting olfactory neurons is unlearnable. **A1.** Paired plot showing the chamber preference index (PI) before and after *Orco* conditioning with a continuous optogenetic-light stimulus. Black dots indicate the mean PI, and the error bars represent 95% CIs. **A2.** Change in chamber preference index (ΔPI) between pre- and post-conditioning is shown. The triangles indicate the individual pre:post ΔPI values; black dots indicate mean ΔPI values with their 95% CI (black line), and the Δ distributions (gray curve). **B1.** Preference index (PI) before and after conditioning with pulsed optogenetic-light is shown. The ends of the blue lines indicate the PI scores per chamber for anti–pulsed-light training. **B2.** Blue triangles indicate changes in preference index (ΔPI) per chamber following training. **C1.** The chamber preference index (PI) before and after conditioning using continuous optogenetic-light stimulus is shown. Mean PI values are denoted by black dots, with 95% confidence intervals shown as error bars. **C2.** Orange triangles mark individual ΔPI values, while black dots represent the mean ΔPI with 95% confidence intervals (black line). The gray curve shows the distribution of ΔPI values. **D1.** The plot shows the preference index (PI) for each chamber before and after conditioning with a pulsed optogenetic-light stimulus. Blue lines connect pre- and post-conditioning PI values for anti–pulsed light training. **D2.** The triangles represent the change in preference index (ΔPI) after training. **E1.** Paired plot presenting the chamber preference index (PI) both before and after conditioning with a continuous optogenetic-light stimulus. Black dots represent the average PI, with error bars indicating the 95% confidence intervals. **E2.** The change in chamber preference index (ΔPI) between pre- and post-conditioning is shown. Individual ΔPI values are represented by triangles, while black dots indicate the mean ΔPI with 95% confidence intervals (black line). The gray curve reflects the ΔPI distribution. **F1.** The chamber preference index (PI) before and after conditioning using a pulsed optogenetic-light stimulus is displayed. Blue lines indicate PI values per chamber for anti–pulsed light training. **F2.** Changes in chamber preference index (ΔPI) after training are depicted by blue triangles.

### Optogenetic activation of dopaminergic neurons implants olfactory memories

After substituting the odor stimulus with optogenetic activation, we investigated whether reinforcing stimuli, either aversive foot shock or rewarding sucrose solution, could be replaced with optogenetic activation. Here, we expected to verify earlier studies showing that punishing reinforcement can be substituted with activity in subsets of TH+ cells that include PPL1s (Claridge-Chang et al. 2009a), and that a sugar reward could be substituted with PAM activation (C. Liu et al. 2012; Burke et al. 2012).

To substitute the foot-shock stimulus, we generated transgenic flies that expressed a blue-light-sensitive channelrhodopsin in a subset of TH+ neurons (*TH-LexA; LexAop-Chr2XXL*) that contain the PPL1 neurons (Berry et al. 2015; Cervantes-Sandoval et al. 2017). These flies were then presented with real odors (MCH or OCT) where one of the odors was paired with a blue-light stimulus (**Figure 4A-C**). We found that the test flies expressing Chr2XXL in the *TH-LexA* subset of TH+ neurons strongly avoided the punished odor after training, as demonstrated by a ΔPI of −0.44 [-0.71, −0.10]. By contrast, the control *LexAop-Chr2XXL* and *TH-LexA* flies displayed little conditioned aversion, with ΔPIs of +0.01 [-0.26, +0.30] and −0.18 [-0.63, +0.15]), respectively (**Figure 4C**).

**Figure 4.**
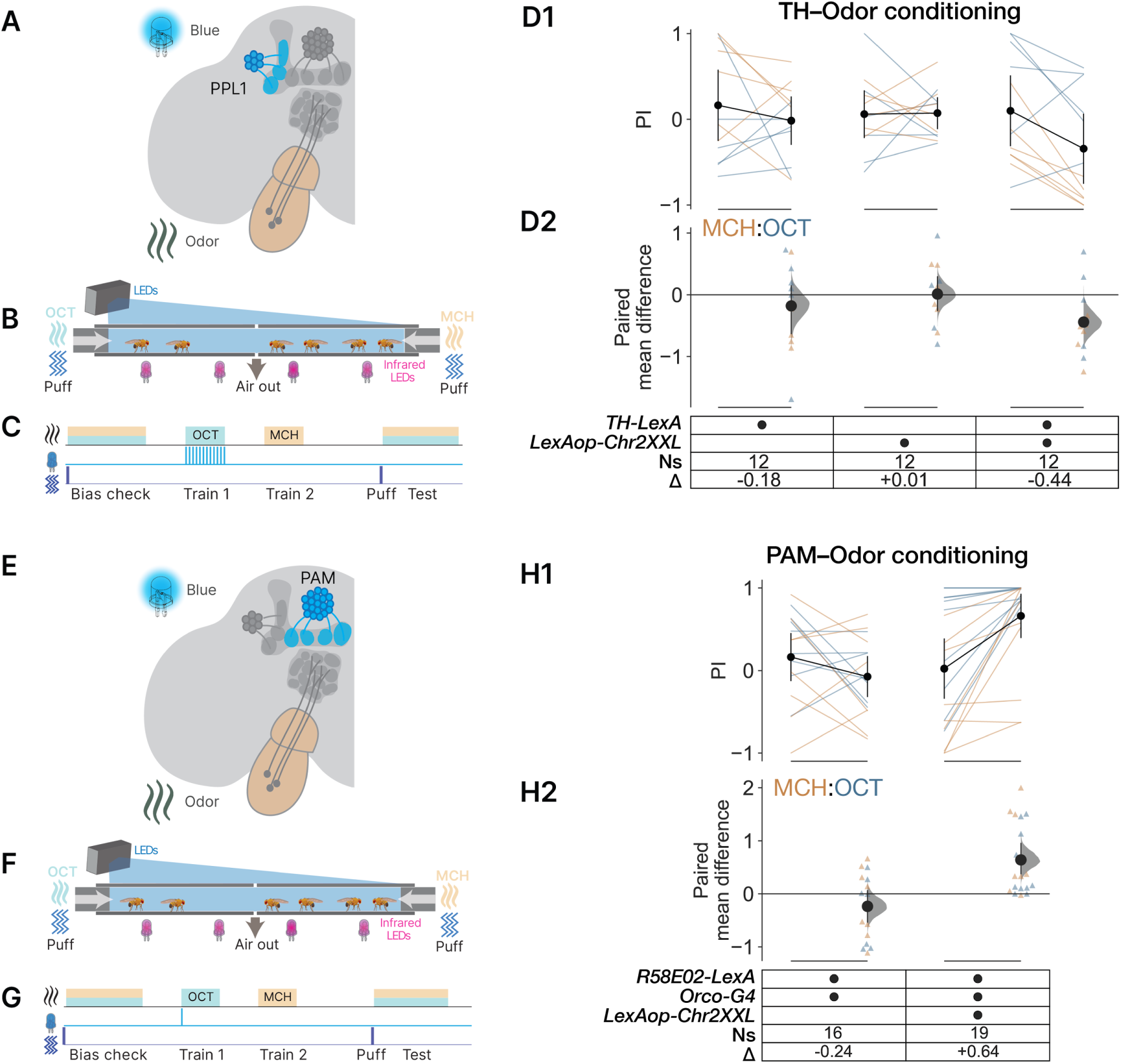
Optogenetic conditioning with activation of *TH-LexA* or *R58E02* neurons induces olfactory memory. **A.** A schematic of the main fly brain regions involved in learning and the stimuli used in the TH-odor conditioning experiments. **B.** Conditioning odors with blue-light activation of ChrXXL expressed in *TH-LexA* cells. **C.** The conditioning protocol: the flies’ naive odor OCT/MCH preference was tested (Bias) before a real odorant was presented along with a blue-light stimulus and applied 12 times (Train 1). A second odor was presented without a light stimulus (Train 2) and then the conditioned odor preference was tested (Test). Both odor regimes were used, but only anti-OCT is depicted. **D1.** Paired plot showing odor preference before and after training. **D2.** Dot plot showing the difference between the pre- and post-conditioning learning scores (represented as ΔPI) per chamber with its mean ΔPI values (black dots) and 95% CI (black line). **E.** Overview of the experimental design to determine if artificial *R58E02* PAM activation can instruct appetitive olfactory-like memories. **F.** Optogenetic conditioning in the MOT assay using light and odor. **G** The timeline of a reward-substitution experiment: the protocol was similar to that used for aversive training, but in Train 1, only a single pulse of blue light was delivered. Only one regime (anti-OCT) is depicted. **H1.** Paired line plot showing the odor preference index (PI) before and after conditioning. **H2.** The change in preference index (ΔPI) after training is shown. Triangles represent the pre:post ΔPI values per chamber, mean ΔPI values (black dots) with its 95% CI (black line), and the Δ distribution (gray curve). Either MCH (orange) or OCT (blue) was used as an odor stimulus in conditioning.

The transgenic flies that expressed Chr2XXL in their *R58E02* PAM neurons similarly developed a preference for odors that were conditioned with blue light, as evidenced by a ΔPI of +0.64 [+0.38, +0.96]. By contrast, the control files exhibited only a modest increase in avoidance (ΔPI = −0.24 [-0.54, +0.06]; **Figure 4E–H**).

### Co-activation of *Orco-Gal4* and *TH-LexA* neurons forms an aversive memory in MOT

To form an aversive associative olfactory memory, two external stimuli are required: in this case, an odor and a foot shock. After showing that the odor and foot shock could be separately substituted with optogenetic activation of the ORNs or *TH-LexA* cells, respectively, we aimed to test if optogenetic activation in both cell types could induce an entirely synthetic memory (**Figure 5A**). To this end, we generated flies that expressed red-actuated Chrimson in *Orco-Gal4* neurons and blue-actuated Chr2XXL in the *TH-LexA* cell cluster, and confirmed the expression of the optogenetic channels by immuno-staining (**Figure 5B**). We then conducted an MOT assay, whereby we subjected flies to a conditioning protocol using both red- and blue-light illumination (**Figure 5C,D**). While control flies exhibited negligible changes in preference, optogenetic flies showed modest conditioned aversion (*TH-LexA/UAS-Chrimson; Orco-Gal4/LexAop-Chr2XXL* ΔPI = −0.33 [-0.64,-0.01], **Figure 5E**). As we saw minimal learning in the control line carrying *TH>Chr2XXL* expression, we concluded that red-light visual learning constitutes a minor component of optogenetic memory (ΔPI −0.08 ≉ −0.33). Of note, the effect size of the fully synthetic memory (−0.33) was smaller than that for both Orco–Shock conditioning (−0.92, **Figure 2D**) and Odor–TH conditioning (−0.44, **Figure 4D**), suggesting that while effective, optogenetic co-activation does not fully recapitulate the shock stimulus.

**Figure 5.**
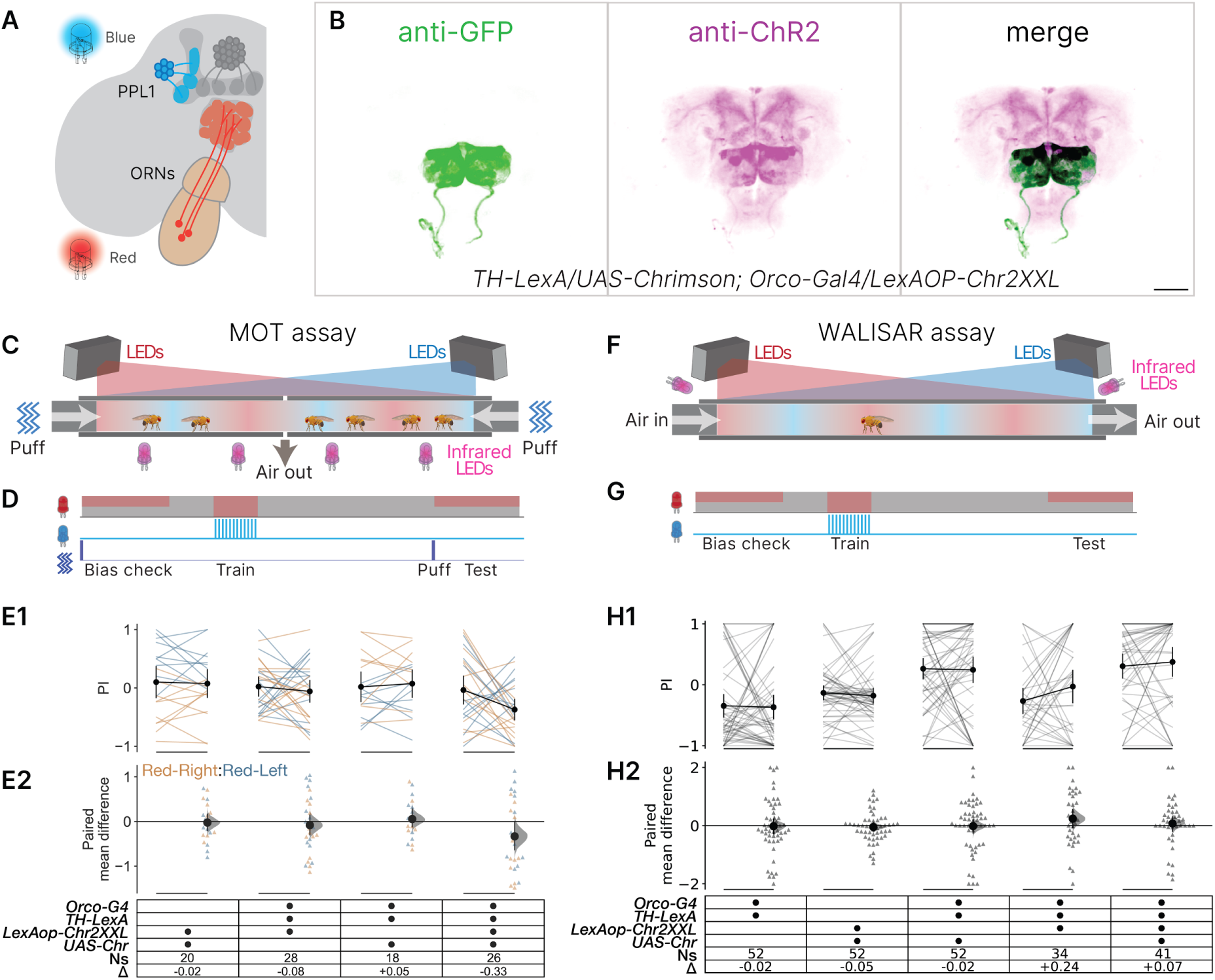
Optogenetic co-activation of *TH-LexA* cells and *Orco-Gal4* neurons can implant aversive memory. **A.** Schematic showing Chr2XXL expression in the *TH-LexA* neurons, and Chrimson expression in the *Orco-Gal4* ORNs. **B.** Confocal images of a two-opsin transgenic brain stained with antibodies against GFP (anti-GFP, reactive with mVenus; green) and channelrhodopsin (anti-ChR, reactive with both ChrXXL and Chrimson; magenta). **C.** Overview of the fully synthetic memory inception experiments with two-opsin optogenetics in the MOT assay. **D.** During the MOT assay, flies were first presented with dark and illuminated domains (Bias); then conditioning with red and blue light simultaneously (Train); lastly, their light–dark preference was re-tested (Test). Red light was used on either the left or right side during training, but only the Red-Left regime is depicted. **E1.** Chamber preference index (PI) before and after conditioning; black dots and error bars represent the mean PI and 95% CI, respectively. **E2.** Orange and blue triangles indicate the individual pre:post ΔPI values for fully synthetic memory inception where either a Red-Right or Red-Left light pattern was used, respectively. **F** Schematic of the WALISAR assay, which uses unidirectional air flow and tracks individual flies. **G** The WALISAR conditioning regimes were identical to those used during the MOT assay, except that there was no air puff at the start of each test. Red light was used either on the left or right, but only the Red-Left regime is depicted. **H1.** Single-fly PI before and after conditioning with blue and red light. **H2.** Individual pre:post ΔPI values per fly are depicted (triangles) with its mean ΔPI (black dot) and 95% CIs (black lines).

We also investigated whether this result could be extended to the simplified WALISAR optogenetic apparatus, which records behavior from individual flies, and has LED illumination with air flow in a single direction (**Figure 5F,G**) (Tumkaya et al. 2022). Here, the same genotype that showed fully optogenetic learning in the MOT assay did not form an aversive olfactory memory in the WALISAR assay (*TH-LexA/UAS-Chrimson; Orco-Gal4/LexAop-Chr2XXL*, ΔPI = +0.07 [-0.20, +0.30], **Figure 5H**). Nevertheless, even though optogenetic aversive conditioning did not generalize to a second apparatus, we conclude that fully optogenetic conditioning is capable of eliciting an aversive memory.

### Co-activation of *Orco-Gal4* and *R58E02* PAM cells drives appetitive memory formation

When an odor is paired with a reward signal that is transmitted to the MB via the dopaminergic *R58E02* PAM neurons, flies will form an appetitive memory (C. Liu et al. 2012; Burke et al. 2012). To test the hypothesis that the coincident optogenetic activation of Orco and PAM neurons will also induce appetitive memory, we generated flies carrying the Chrimson and Chr2XXL channels in these cell types, respectively (**Figure 6A**). We first confirmed the expression of the opsins by immunostaining (**Figure 6B**). Then, we activated the Orco neurons together with the reward-encoding PAM neurons in both groups of flies and individual flies, using the MOT and WALISAR assays, respectively (**Figure 6C–H**). Flies with Chrimson in the *Orco-Gal4* neurons and Chr2XXL in the *R58E02-LexA* PAM neurons and that were conditioned in groups in the MOT assay, developed a preference for Orco-neuron activation by red light, with a ΔPI of +0.50 [+0.24, +0.74] (**Figure 6E-F**). Similarly, optogenetic flies individually trained in the WALISAR assay also developed a preference for Orco-neuron activation, with a ΔPI of +0.38 [+0.10, +0.63] (**Figure 6C-D**). In both experiments, visual learning was a minor component as the *R58E02>Chr2XXL* controls underwent only trivial red-light learning. We thus conclude that synthetic learning can be induced with *R58E02* + Orco activation, and is comparable with *R58E02*-induced olfactory learning (+0.50 *vs* +0.64, Figure 6E, 2H) and is robust across two assay formats.

**Figure 6.**
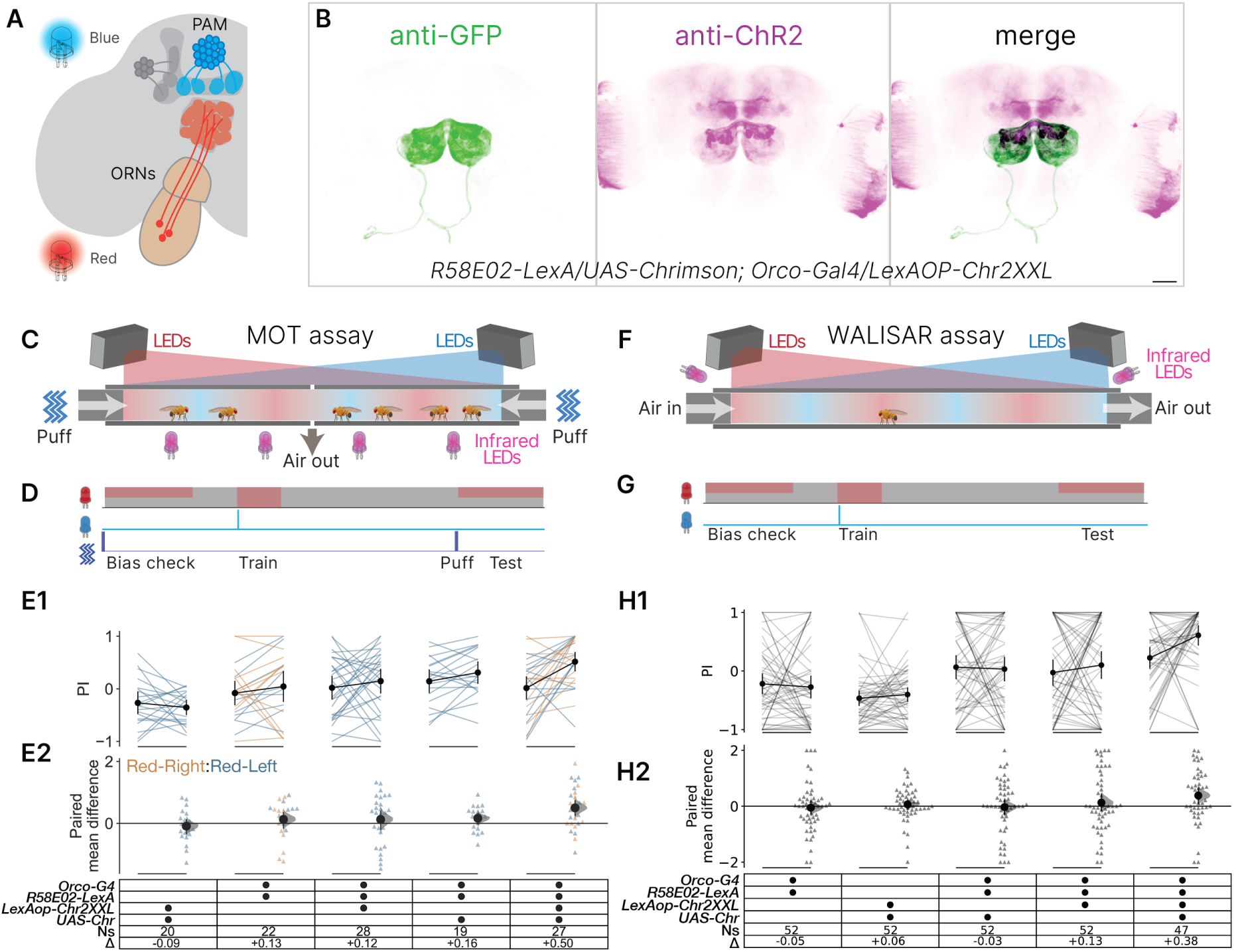
Optogenetic co-activation of Orco and PAM cells elicits appetitive memory. **A** Schematic showing Chr2XXL expression in *R58E02-LexA* PAM neurons and Chrimson in ORNs. **B** Confocal images of a transgenic brain expressing opsins driven by *R58E02-LexA* and *Orco-Gal4*. Anti-ChR2 stained both Chr2XXL and Chrimson (magenta) while anti-GFP stained the mVenus tag on Chrimson (green). **C** Schematic of the optogenetic conditioning protocol via MOT assay. **D** The MOT conditioning protocol, where flies were tested in groups for their red-light preference (Bias), then subjected to red light and a single pulse of blue light (Conditioning) before their red-light preference was tested again (Test). **E1.** Red-light preferences before and after conditioning in the MOT assay. Two different timings of the training epoch were used. **E2.** Dot plot showing the pre:post ΔPI per chamber in fully artificial appetitive memory conditioning experiments. Either a Red-Right (orange) or Red-Left (blue) light pattern was used in the test. Black dots indicate the mean ΔPI, and the error bars indicate the 95% CIs. **F.** Schematic of the optogenetic conditioning protocol via WALISAR assay. **G.** The WALISAR conditioning regime was established as described in D for the MOT assay. **H1.** Red-light preferences of individual flies before and after light conditioning. **H2.** The change in preference index (ΔPI) of individual flies before and after training is shown. Gray triangles indicate the pre:post ΔPI values per fly in fully synthetic memory implantation experiments in WALISAR. Black dots indicate mean ΔPI values, and the error bars indicate the 95% CIs.

## Discussion

### Verification of semi-synthetic conditioning

In this study, we aimed to test the efficacy of conditioning the olfactory system with optogenetic co-activation in order to further dissect how olfactory signals are encoded by the *Drosophila* brain. To do so, we constructed experimental paradigms that enabled the control of odor, shock, and two light sources, and used them to test stimulus-evoked and optogenetically incepted memories. By this approach, we found that pairing foot shocks with optogenetic activation of ORNs led to the formation of aversive memories. Similarly, synthetic activation of *TH-LexA* or *R58E02* PAM neurons during exposure to odorants led to robust preference changes in the averted or attracted direction. Our optogenetic conditioning protocol was originally modeled after the conventional olfactory-memory shock-conditioning protocol, which typically delivers 12 foot shock pulses over one minute (1.5 s ON – 3.5 s OFF) (Tumkaya, Ott, and Claridge-Chang 2018). While this protocol immediately yielded a robust inception for aversive conditioning, no inception was observed for appetitive conditioning. The single-pulse protocol, however, did yield robust appetitive memory inception. These results indicate that the long Chr2XXL opening time does not affect aversive conditioning, but may be the reason that 12-pulse appetitive conditioning was ineffective, and may identify a difference between the PPL1 and PAM systems (Dawydow et al. 2014). Overall, the effect sizes in the three optogenetic paradigms were comparable with natural-stimulus conditioning, establishing that optogenetic activation can substitute for sensory inputs to induce memories in semi-artificial learning paradigms.

### Implanting fully fictive memories

Dual optogenetic activation of Orco neurons and dopaminergic cells (*TH-LexA* or *R58E02* PAM neurons) induced robust memory inception in three of four experiments. Comparing effect sizes across aversive MOT conditioning experiments, the double-optogenetic memory was ΔPI = –0.33, slightly smaller than Odor+TH memory (–0.44), around half the size of natural odor+shock memory (–0.59 average), and markedly smaller than shock+Orco memory (–0.92). Similarly, in MOT attraction conditioning, the fully synthetic memory was slightly smaller than the semi-synthetic memory (0.50 < 0.64). In the WALISAR single-fly assay, both *TH-LexA*+*Orco* and *R58E02*+*Orco* fully synthetic memories were reduced relative to MOT effect sizes (–0.07 > –0.33 and 0.38 < 0.50). We posit that this trend of smaller effect sizes in WALISAR could stem from sampling error, or differences from the MOT assay, such as the design of the apparatus, use of an air puff at the start of each trial, and social cues from conspecifics during the trial in the MOT assay (Muria et al. 2021).

While it is true that CO_2_ released by groups of flies can have an enhancing effect on olfactory memory (Muria et al. 2021), the success of single-fly aversive conditioning in a previous study demonstrates that it is not essential for robust memory formation (Claridge-Chang et al. 2009b). Future investigations systematically comparing single versus group of flies are needed to elucidate the group effect on olfactory learning. Nevertheless, for *R58E02* PAM conditioning, the flies became attracted to Orco-neuron activation in both assays, indicating the robustness of appetitive memories.

### Implications for memory formation

Fully synthetic memory implantation has already been demonstrated in two systems: *Drosophila* larvae and mice (Honda et al. 2014; Vetere et al. 2019). Like the present study, the experiments conducted in these model systems used heterologous activation of olfactory receptors. The larval experiments conditioned Or24a neurons with the activation of Tyrosine decarboxylase 2 (Tdc2) cells, which produce octopamine and tyramine neuromodulators (Honda et al. 2014). The mouse study conditioned *Olfr160*/M72-bearing olfactory sensory neurons with the co-activation of two distinct projections into the ventral tegmental nucleus. Memory was tested by showing the association with a known odorant ligand of M72, acetophenone, establishing transferability between synthetic and natural stimuli. While it has been claimed that fully synthetic cell type-specific optogenetic conditioning directly supports (Vetere et al. 2019; Josselyn and Tonegawa 2020) the synaptic-plasticity hypothesis of memory formation (Martin and Morris 2002), these experiments do not manipulate synapses directly, so do not speak to hypotheses of synaptic-weight changes *per se*.

Our study extends these findings by showing that ORN activation can be conditioned with coincident dopaminergic activation. These findings provide a definitive answer to a fundamental inquiry in neuroscience: artificial, rectangular-pulse sensory and neuromodulatory stimuli can instruct associative memory formation.

According to the classical rate-coding theory (Adrian and Zotterman 1926; Stein, Gossen, and Jones 2005), information is represented in the brain by the number of neuronal spikes during a stimulus epoch. Temporal-coding theory (Buonviso, Cenier, and David 2009; Menini 2009), on the other hand, holds that firing rate itself is insufficient, and that the precise timing of neuronal spikes is instrumental in encoding stimulus features and has a vital role in odor perception, as well as learning and memory (Perl et al. 2020; Stern et al. 2018; Dragoi 2020; Lin, Huang, and Richards 2023). Neuronal firing patterns of the fly olfactory neurons upon artificial activation using channelrhodopsins have been recorded by several groups (Hallem, Ho, and Carlson 2004; Bellmann et al. 2010; Inagaki et al. 2014; Klapoetke et al. 2014; Bell and Wilson 2016; Fox and Nagel 2021; Zanon, Zanini, and Haase 2022). These studies show that, while not square *per se*, the firing patterns elicited by rectangular-pulse optogenetic stimuli are largely unstructured and do not mimic the temporal patterns of natural stimuli. The success of the fully synthetic experiments in the present study shows that (in the neurons that were targeted) rate coding is instructive, as the data show that associative memory does not have any strict requirements for temporal or ensemble patterns of activity in ORNs and DANs that are specific to natural stimuli (e.g. odorants, sugar, or pain); rather, it verifies that simple activity coincidence in these systems is sufficient as an effective driver of memory formation.

### Study limitations

An important caveat to our study is that ORNs themselves are not the primary site of memory formation; rather, they transmit odor information via at least two serial synapses to the MBs, where the dopaminergic inputs are thought to act on synapses formed by the intrinsic MB neurons, Kenyon cells. The signal from the simple activation of all *Orco-Gal4* ORNs is likely transformed in downstream neurons, gaining temporal structure, and eliciting additional ensemble patterns (Wilson, Turner, and Laurent 2004; Wilson 2013; Gruntman and Turner 2013; Groschner and Miesenböck 2019; Benton 2022). Thus, while our findings conclusively demonstrate that temporal structure is dispensable to dopaminergic neurons and ORNs for memory formation, its relevance to higher-level olfactory encoding requires further investigation. Future experiments substituting the odor stimulus via optogenetic activation of Kenyon cells (KCs), rather than ORNs, could address this question. The success of a previous study demonstrating that thermogenetic activation of random and sparse KC ensembles can serve as a CS in a semi-artificial learning paradigm (Vasmer et al. 2014) suggests that dual-optogenetic activation of the dopaminergic neurons and KCs would form memories.

### Implications for future studies

Fully optogenetic conditioning enables novel paradigms for interventional memory experiments, where precise temporal and cellular control of activity is crucial for delineating memory mechanisms, such as contingency protocols, operant paradigms, trial repetition, retention, and extinction. For instance, researchers have previously investigated the impact of the interstimulus interval on memory formation, defined as the temporal gap between the onset of the CS and US inputs, and discovered that synaptic strength is highly sensitive to the millisecond-scale temporal correlation between the input signals. (Cassenaer and Laurent 2012; Martinez-Cervantes et al. 2022; Kjell, Löwgren, and Rasmussen 2018; Akins 2019; Zhao et al. 2021). Interestingly, growing evidence from animal models suggests that forming associations between temporally separated stimuli, rather than contiguous ones, invokes distinct molecular pathways, and may also require higher-order cognitive functions that have evolved from the capacity for contiguous stimuli learning, such as attention and awareness (Dylla et al. 2013; Shuai et al. 2011; Paoli, Macri, and Giurfa 2023). As dual-optogenetic conditioning enables the precise temporal control of CS and US signals, it could allow investigations into such questions about US–CS delays, or fine temporal structure of activity. This, in turn, could reveal how neural systems have evolved to store memories, and thereby guide new research directions and translational work on memory disorders.

In summary, our findings demonstrate that fully synthetic optogenetic inception is generalizable to dopaminergic neurons in adult *Drosophila*. These findings establish that simple co-activation with ORNs is sufficient for both aversive and attractive memory formation. Moving forward, leveraging the fully optogenetic approach for investigating underlying circuits stands to substantially improve our understanding of memory dynamics.

## Acknowledgments

The authors would like to thank Christian Wegener (University of Wurzburg) for providing the *LexAop-Chr2XXL* flies, and Dr Ravinuthula Sruthi Jagannathan (Duke-NUS Medical School) for their critical review of the manuscript before submission.

## Author Contributions

Conceptualization: TT, ACC; Experiment design: TT, XY, ACC; Methodology: TT, XY, ACC; Software: TT, YM (Python); Data Analysis: TT, YM (Python), JCS (CRITTA, LabView); Investigation: TT (genetics, fly husbandry, behavior, immunohistochemistry, and microscopy), XY (genetics, behavior, brain dissection, immunohistochemistry, microscopy); Resources: JCS (instrumentation); Writing – Original Draft: TT; Writing – Revision: TT, XY, YM, ACC; Visualization: TT, YM, XY, ACC; Supervision: ACC; Project Administration: ACC; Funding Acquisition: ACC.

## Competing interests

The authors declare no competing interests.

## Funding

TT, YM, XYZ, and ACC were supported by grants MOE2017-T2-1-089, MOE2019-T2-1-133, and MOE-T2EP30222-0018 from the Ministry of Education, Singapore; JCS and ACC were supported by grants 1231AFG030 and 1431AFG120 from the A*STAR Joint Council Office. YM was supported by Duke-NUS Medical School and by a President’s Graduate Fellowship funded through the Jasmine Scholarship and MOE-T2EP30222-0018 (Research Scholarship); XYZ was supported by a Yong Loo Lin School of Medicine scholarship and FY2022-MOET1-0001. The authors were supported by a Biomedical Research Council block grant to the Institute of Molecular and Cell Biology, and a Duke-NUS Medical School grant to ACC.

## Notes

https://zenodo.org/records/13935195

### Competing Interest Statement

The authors have declared no competing interest.

